# Impact of changes in buffer ionic concentration and mutations on a GH1 β-Glucosidase homodimer

**DOI:** 10.1101/2025.02.10.637493

**Authors:** Rafael S. Chagas, Sandro R. Marana

## Abstract

Oligomerization is a key feature of protein function, with approximately 30% of proteins exhibiting this trait. The homodimeric form of proteins, such as the GH1 β-Glucosidase from *Spodoptera frugiperda* (Sfβgly), plays a significant role in enzyme activity. In this study, we investigate the homodimerization of Sfβgly, which forms a cyclic C2 dimer with a well-defined interface. Using size exclusion chromatography and SEC-MALS, we characterized the homodimerization behavior of Sfβgly at equilibrium conditions in different ionic concentrations of phosphate buffer. The dissociation constants (*K*_D_) increase with decreasing ionic concentration, suggesting that the hydrophobic effect is central to homodimer formation. Site-directed mutagenesis of key residues at the dimer interface further elucidated the contributions of specific amino acid residues to dimer stability. Mutations affecting both, apolar and hydrogen bond-forming residues, significantly increased the *K*_D_. However, mutations of hydrogen bond-forming residues caused a smaller *K*_D_ change than apolar residue mutations, suggesting that while the latter is the driving factor in the dimerization, the former may play a crucial role in guiding the monomers relative orientation. These findings enhance our understanding of protein oligomerization in GH1 β-Glucosidases and its implications for protein design and function.

## 1. Introduction

Oligomerization is a well-documented phenomenon in proteins, crucial for their biological function. This process occurs in all the life forms and involves approximately 30% of all the proteins [1, 2]. Several significant proteins such as enzymes active upon DNA [3], tyrosine kinase receptors [4], G-protein-coupled receptors [5], transcriptional factors [6], and ion channels [7] are oligomers.

Oligomeric proteins are composed of multiple subunits (polypeptide chains), which may be identical (a homo-oligomeric protein) or different (hetero-oligomers) [8]. The term homomer has been commonly used instead of homo-oligomer to refer to a protein oligomer formed by self-interacting copies of a protein subunit [1]. The most frequent forms of homomers are dimers and tetramers, which occur approximately four times more often than hetero-oligomers [9].

The formation of homomers provides several structural and functional benefits [10], including enhanced stability [11, 12], control over the accessibility and specificity of active sites [3], and structural complexity. It also minimizes genome size [13]. After all, when a protein exists as a dimer, it requires only a single encoding in the genome, unlike a monomeric protein of the same molecular weight. This concept is referred to as “genetic saving” [14].

In the UniProt Database, the most comprehensive protein database, homodimeric proteins make up over 51%, significantly surpassing the proportion of monomeric and other oligomeric forms [14]. According to the BRENDA enzyme database [15], the majority of enzymes are homomers, with dimers being more prevalent than monomeric enzymes [15].

Here, we investigated the homodimer of the GH1 β-Glucosidase from *Spodoptera frugiperda* (Sfβgly), a secreted digestive enzyme associated with the glycocalyx of midgut epithelial cells in the cited armyworm [17]. This enzyme has been extensively studied regarding its catalysis, thermal stability, substrate recognition, and oligomerization [17, 18, 19, 20]. General features for GH1 β-Glucosidases, enzymes grouped based on their sequence [21], include the TIM barrel ([β/α]_8_ barrel) fold, the hydrolysis of O- or S-glycosidic linkages, and a tendency to form homomers. However, the regions involved in the formation of the oligomerization interface vary from one enzyme to the other [22], reflecting the diversity of their quaternary structures [23].

Sfβgly (5CG0 PDB code) exhibits a cyclic symmetry, specifically a C2 homodimer, which means that rotating the complex by 180° around the symmetry axis results in the subunits being mutually superposed [24]. In this type of symmetry, the contact between subunits is made by the same surface in both monomers, which is referred to as an isologous interface [25]. The dimeric interface in Sfβgly encompasses 905 Å^2^, or 5% of the total surface area in each monomer. It is composed of 30 residues, 63% of which are apolar, and 4 residues in each monomer are involved in hydrogen-bond formation. The interface presents no disulfide bond or salt bridges (Figure 1, Table 1).

**Figure 1.**
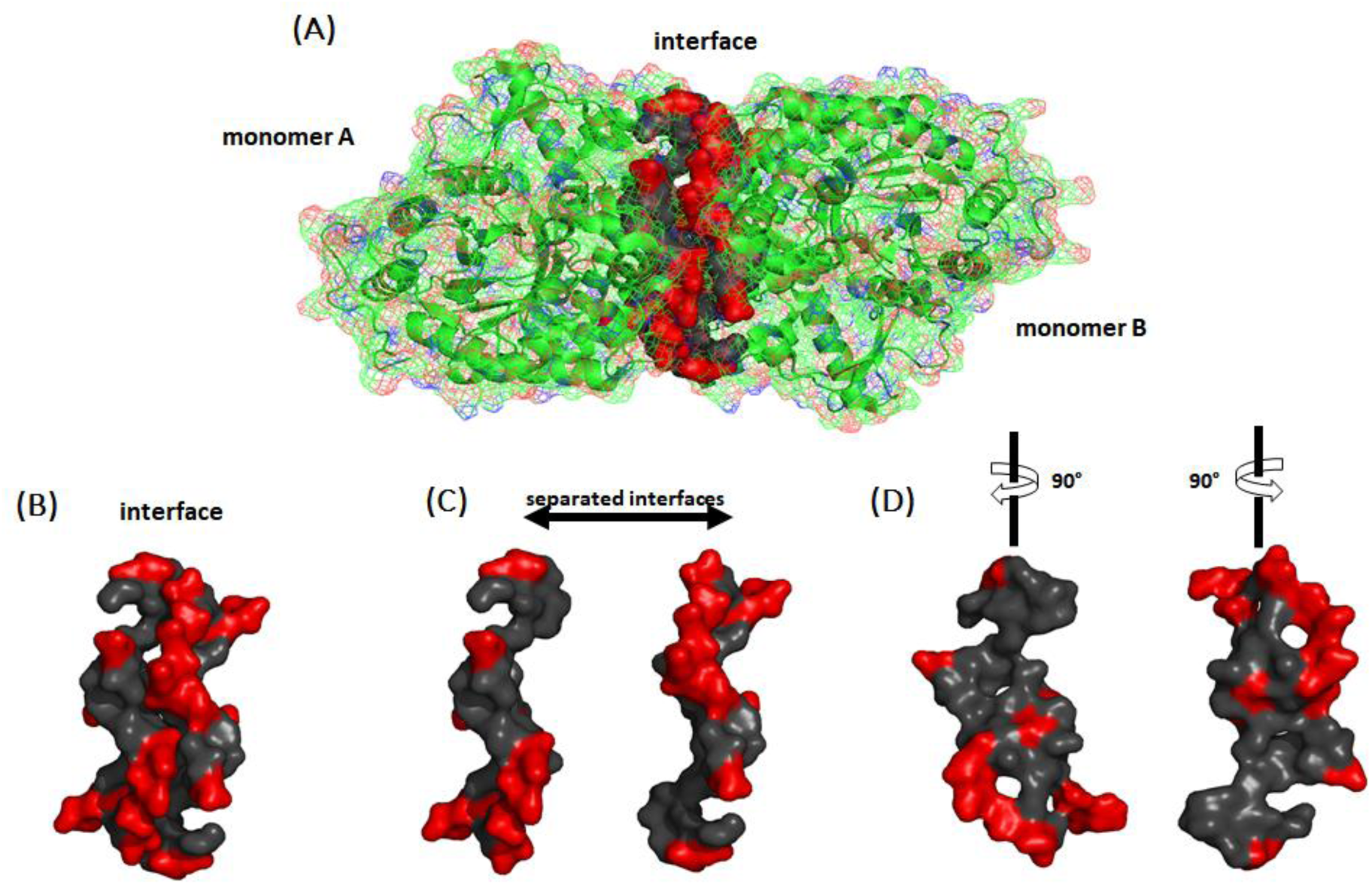
Representation of the Sfβgly homodimer (PDB 5CG0). (**A**) Interface residues are represented as a solid molecular surface (grey and red), whereas the two monomers are shown as cartoons covered with a wired superficial envelope. Apolar residues are in gray and polar residues are in red. (**B**) Dimerization interface surface. (**C**) Separated dimerization interface surface. (**D** oth surfaces were rotated. Their internal composition is predominantly apolar (grey).

**Table 1.** Residues forming the Sfβgly homodimer interface.

| Residue | Surface area buried in the interface (%) | Residue | Surface area buried in the interface (%) |
| --- | --- | --- | --- |
| T108 | 10 | R169 | 10 |
| M110 | 80 | Y196 | 20 |
| A111 | 70 | L205 | 70 |
| N112 | 80 | N206 | 10 |
| K148 | 10 | A207 | 40 |
| Q150 | 20 | M210 | 70 |
| E151 | 60 | G211 | 80 |
| L152 | 90 | L214 | 70 |
| G153 | 100 | N218 | 10 |
| A156* | 100 | Q301 | 30 |
| N157* | 100 | G302 | 40 |
| P158 | 60 | Y303 | 80 |
| L159 | 60 | P304 | 30 |
| W163 | 60 | W305* | 40 |
| D166* | 80 | P348 | 10 |

The lines occupied by apolar residues are marked in gray. Residues involved in hydrogen bonds are marked with an asterisk. Interface residues were identified by using the PDBePISA based on the crystallographic structure 5CG0.

Oligomerization in GH1 β-Glucosidases has been hypothesized as an adaptive strategy for thermal stability [26, 27]. Also, it was demonstrated for Sfβgly that oligomerization turns the homodimer more active than the monomer, and that the oligomerization occurs through a conformational selection mechanism, *i.e.* monomers exchange between two conformations in equilibrium and only one of them combine with a similar one forming the homodimer [20].

The phenomenon of protein oligomerization can be modulated by some factors. Natural modulators include protein concentration, ionic strength, pH, effectors, and nucleic acids while examples of artificial modulators are cross-linkers, shiftides, and nanoparticles [2]. In the present study, we investigated the Sfβgly homodimer formation in a range of different concentrations of phosphate buffer, which made it possible for us to estimate the homodimer dissociation constant (*K*_D_) in different ionic strengths. We also introduced a series of mutations directed to the homodimer interface, permitting us to determine the individual contributions of those residues to the homodimerization. The results showed that the hydrophobic effect is the main component that leads to dimerization in Sfβgly, whereas residues involved in hydrogen bond formation probably act as guidance to the monomers relative orientation.

## 2. Results

### 2.1 Characterizing the dimerization of Sfβgly in different ionic concentration buffers

The wild-type Sfβgly was purified and its purity was verified using SDS-PAGE (Supplementary Figure 1). Its oligomeric states were assessed through size exclusion chromatography (SEC) in different concentrations of the sodium phosphate buffer. Sfβgly appeared as both a monomer (retention volume ≈ 25 mL) and a dimer (retention volume ≈ 30 mL) in 100 mM sodium phosphate buffer pH 6 (P100). Their identity was assessed and confirmed by SEC-MALS (Supplementary Figure 2). The same oligomeric states were observed in 75 mM, 50 mM, 25 mM, and 5 mM sodium phosphate buffers pH 6 (P75, P50, P25, and P5, respectively) (Figure 2).

**Figure 2.**
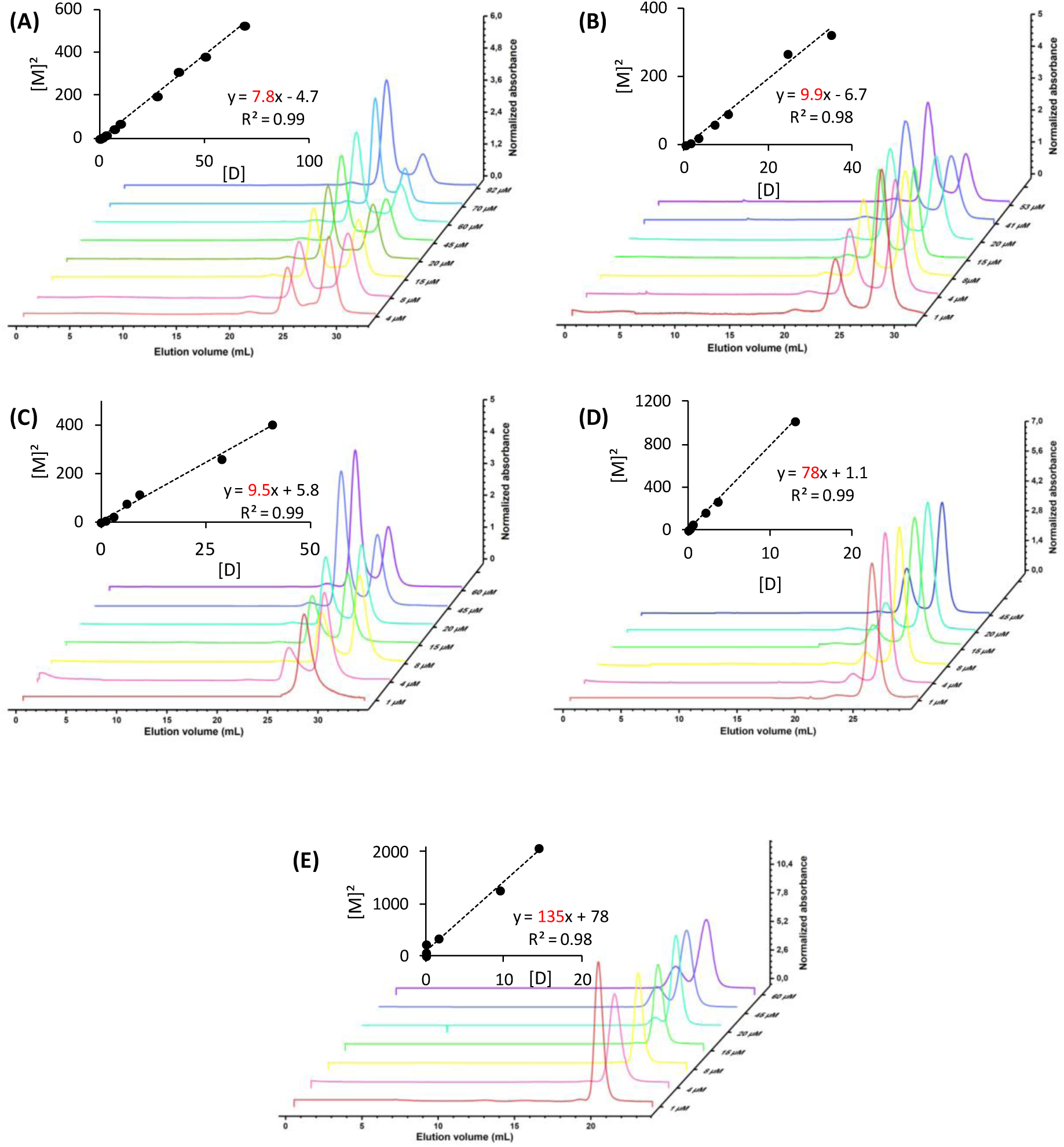
Estimation of the *K*_D_ of the Sfβgly homodimer at different ionic concentrations. (**A**) P100; (**B**) P75; (**C**) P50; (**D**) P25; (**E**) P5. [M] stands for monomer concentration. [D] is homodimer concentration. Line slopes, which correspond to the *K*_D_, are shown in red. Experiments were conducted at pH 6 at 5°C. See Materials and Methods for details.

Across all buffer concentrations tested, the balance between the homodimer and monomer changed as different concentrations of Sfβgly were examined (Figure 2). In P100, with higher Sfβgly concentrations, the dimeric state predominated. But, as the Sfβgly concentration decreased, the balance gradually reversed, with the monomeric state becoming more prevalent. The same pattern was observed in the other buffer concentrations. However, as the buffer concentration decreased, a higher concentration of Sfβgly was required to shift the equilibrium into the dimer prevalence. That reflects changes in the dimer dissociation constant (*K*_D_).

To assess the effect of the ionic concentration on the dimerization process, we determined the *K*_D_ based on the area of the chromatographic peaks (Figure 2) by assuming a simple dimerization equilibrium, M + M ⇌ D.

Thus, the homodimer *K*_D_ remained stable at P100, P75, and P50, *K*_D_ = 7 ± 1, 9.9 and 9.5 µM, respectively, but it presented more pronounced changes at P25 and P5, *K*_D_ = 78 and 135 µM, respectively (Figure 3).

**Figure 3.**
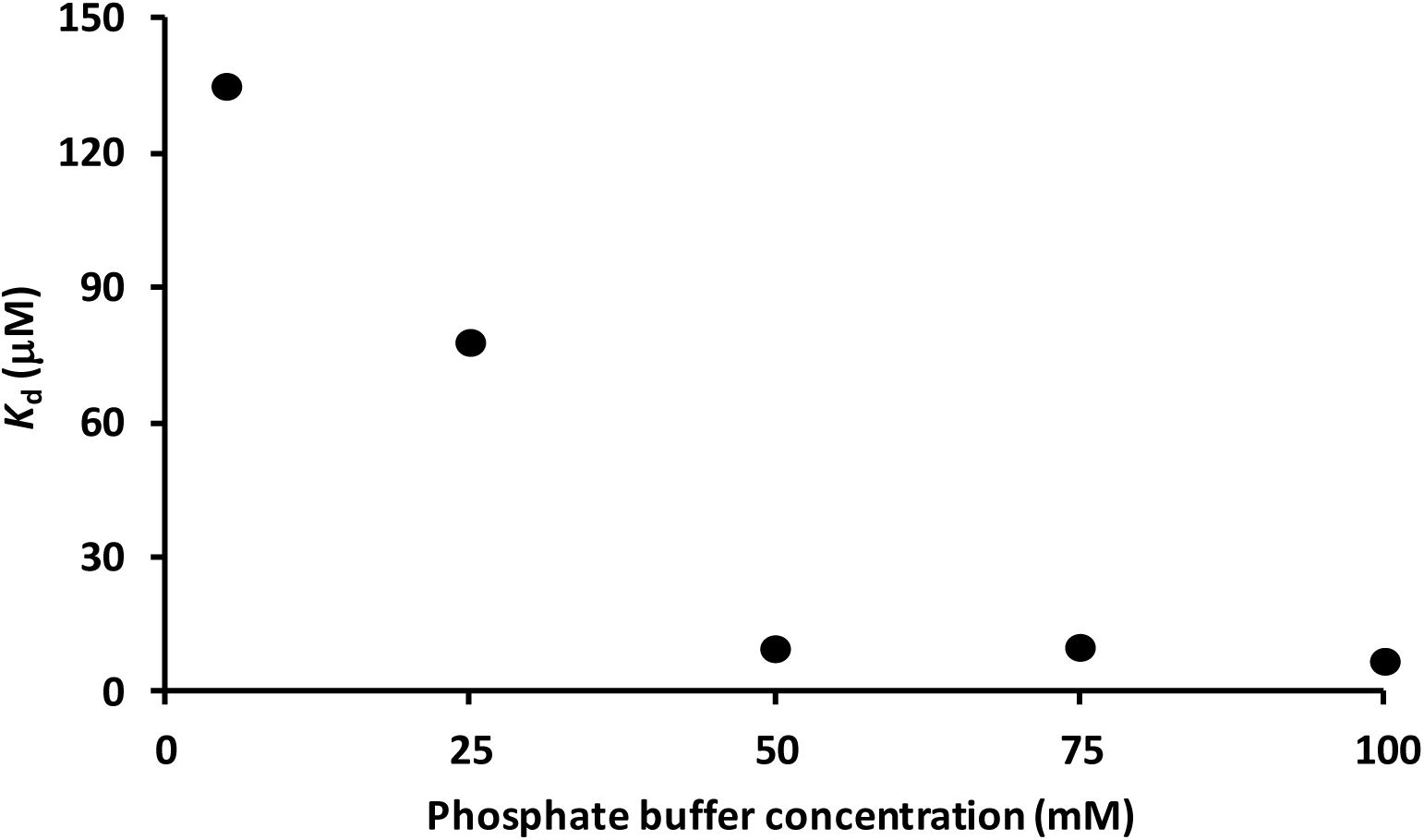
Effect of the phosphate buffer concentration on the dissociation constant (*K*_D_) of the Sfβgly dimer.

In summary, the increment in the concentration of the phosphate buffer resulted in a starkingly decrease of the *K*_D_ (Figure 3), indicating a drastic increase in the affinity between the Sfβgly monomers. Just as a comparison, a similar effect is observed in hydrophobic interaction chromatographies. Conversely, if the monomer binding were mainly driven by non-covalent interactions based on electric charges (ionic bridges and hydrogen bonds), the rise in the ionic concentration would hamper the dimerization, *i.e.* reduce the *K*_D_. Hence, Figure 3 suggests that the hydrophobic effect is the main component impeling the Sfβgly dimerization.

### 2.3 Detailing the dimerization of mutant Sfβgly

Previous experiments composed a broad perspective of the forces guiding the Sfβgly dimerization. However, individual residues of the dimerization interface may have different relative contributions to that process. Thus, point mutations were introduced at dimeric residues presenting more than 70% of their surface area buried in the interface. Thus, six different mutant Sfβgly, each of them with a single replacement (N112S, N157S, D166S, M210A, L214A, and Y303A), were expressed as folded soluble proteins in bacteria and purified (Supplementary Figure 1; Supplementary Figure 4).

The *K*_D_ of the mutant Sfβgly homodimers were evaluated using the same methodology employed for the wild-type Sfβgly (Figure 4, and Table 2).

**Figure 4.**
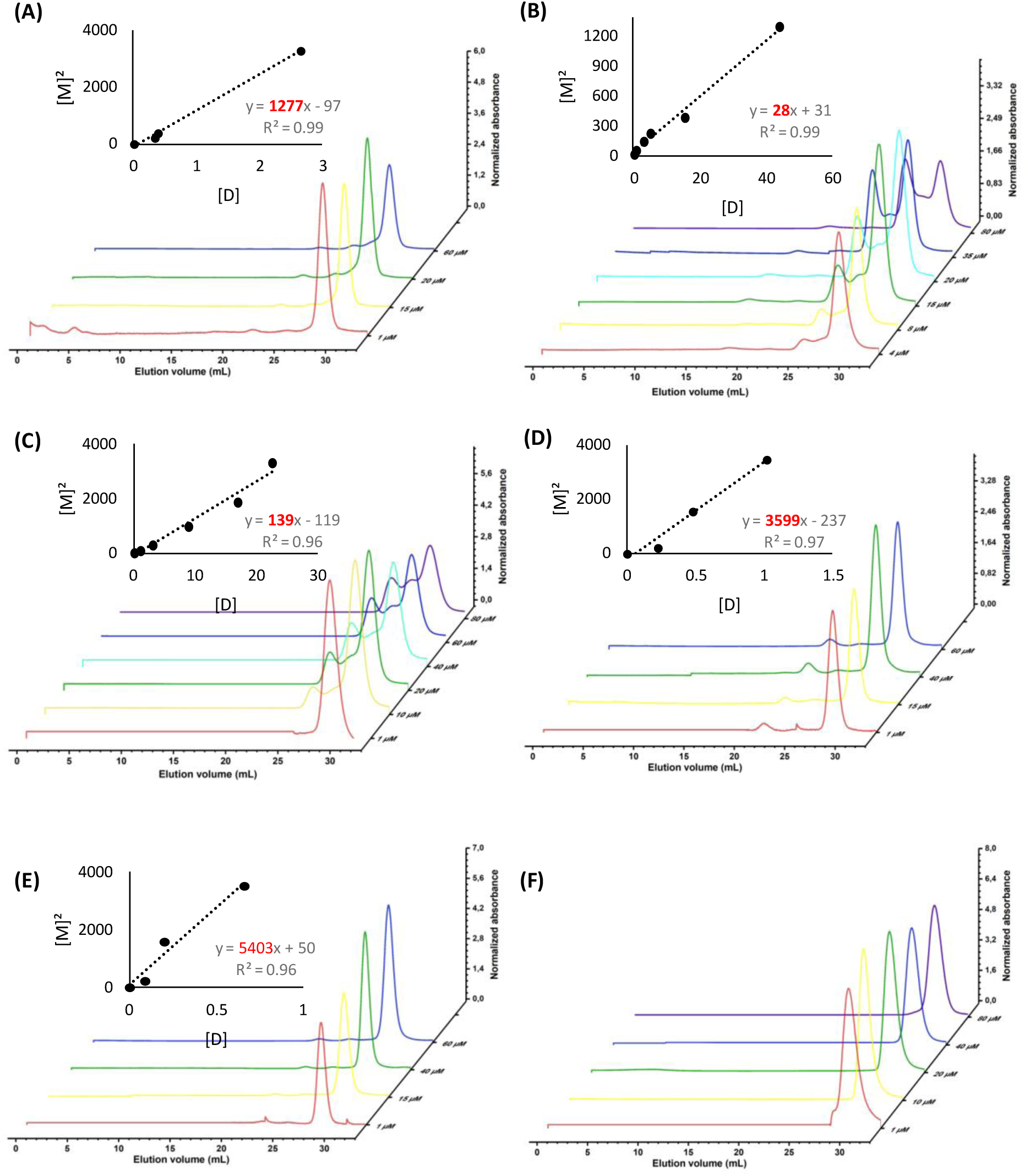
Estimation of the *K*_D_ of the mutant Sfβgly homodimer in P100. **(A)** N112S, **(B)** N157S*, **(C)** D166S*, **(D)** M210A, **(E)** L214A and **(F)** Y303A. Mutant Y303A was observed as a monomer. Asterisks indicate mutations of residues involved in hydrogen bond between the monomers in the interface of the wild-type protein dimer.

**Table 2.** The *K*_D_ and free energy change (ΔΔ*G*^0^_diss_) of the homodimer dissociation for the wild-type and mutant Sfβgly.

| Sfβgly | $K_D$ (mM) | $\Delta\Delta G_{\text{diss}}^0$ (kJ/mol) |
| --- | --- | --- |
| wild-type | 0.008 | - |
| N112S | 1.2 | 13 |
| N157S* | 0.028 | 5.9 |
| D166S* | 0.1 | 2.4 |
| M210A | 3.6 | 15 |
| L214A | 5.4 | 16 |
| Y303A | nd | nd |

nd, not determinable. Mutant Y303A was observed as a monomer. So, for mutant Y303A the $K_D$ and $\Delta\Delta G_{\text{diss}}^0$ must be higher than the highest observed, *i.e.* > 5.4 mM and > 16 kJ/mol. Asterisks indicate mutations of residues involved in hydrogen bond between the monomers in the interface of the wild-type homodimer.

#### 2.3.1 Residues involved in the hydrophobic effect

The single mutations N112S, M210A, and L214A were sufficient to drastically increase the *K*_D_ of the dimer, causing an increase of three orders of magnitude. Mutant Y303A was found only in the monomeric form, suggesting an even more drastic increase in the *K*_D_. Using the relationship between *K*_D_ and Δ*G*^0^_dissociation_, ΔΔ*G*^0^_diss_ = *RT* Ln (*K*_D wild-type_ / *K*_D mutant_), the average loss in affinity between the monomers is approximately 15 *k*J/mol (Table 2), which corresponds to about 50% of the binding energy between the monomers (29 *k*J/mol; based on the *K*_D_ of the wild-type Sfβgly dimer). The loss of affinity resulting from the Y303A mutation must be even more significant, certainly surpassing the above values and preventing dimer formation.

#### 2.3.2 Residues involved in the hydrogen bonds

Moving forward, we also tested the N157S and D166S mutants, which allowed an estimation of the hydrogen bond energy at the interface of the Sfβgly homodimer. There are four hydrogen bonds between the monomers in the dimer, N157 acting as a donor in the interaction with A156 and D166 as an acceptor in the interaction with W305 (Figure 5). Thus, each monomer has 2 donors and 2 acceptors involved in the formation of the four hydrogen bonds. Therefore, just two mutations, N157S and D166S, are enough to evaluate the role of the four hydrogen bonds at the homodimer interface. Notably, there is one hydrogen bond at each edge of the interface and two in the center, with the spatial distribution of the donors and acceptors allowing only one relative orientation of the monomers where all four bonds are formed (Figure 5).

**Figure 5.**
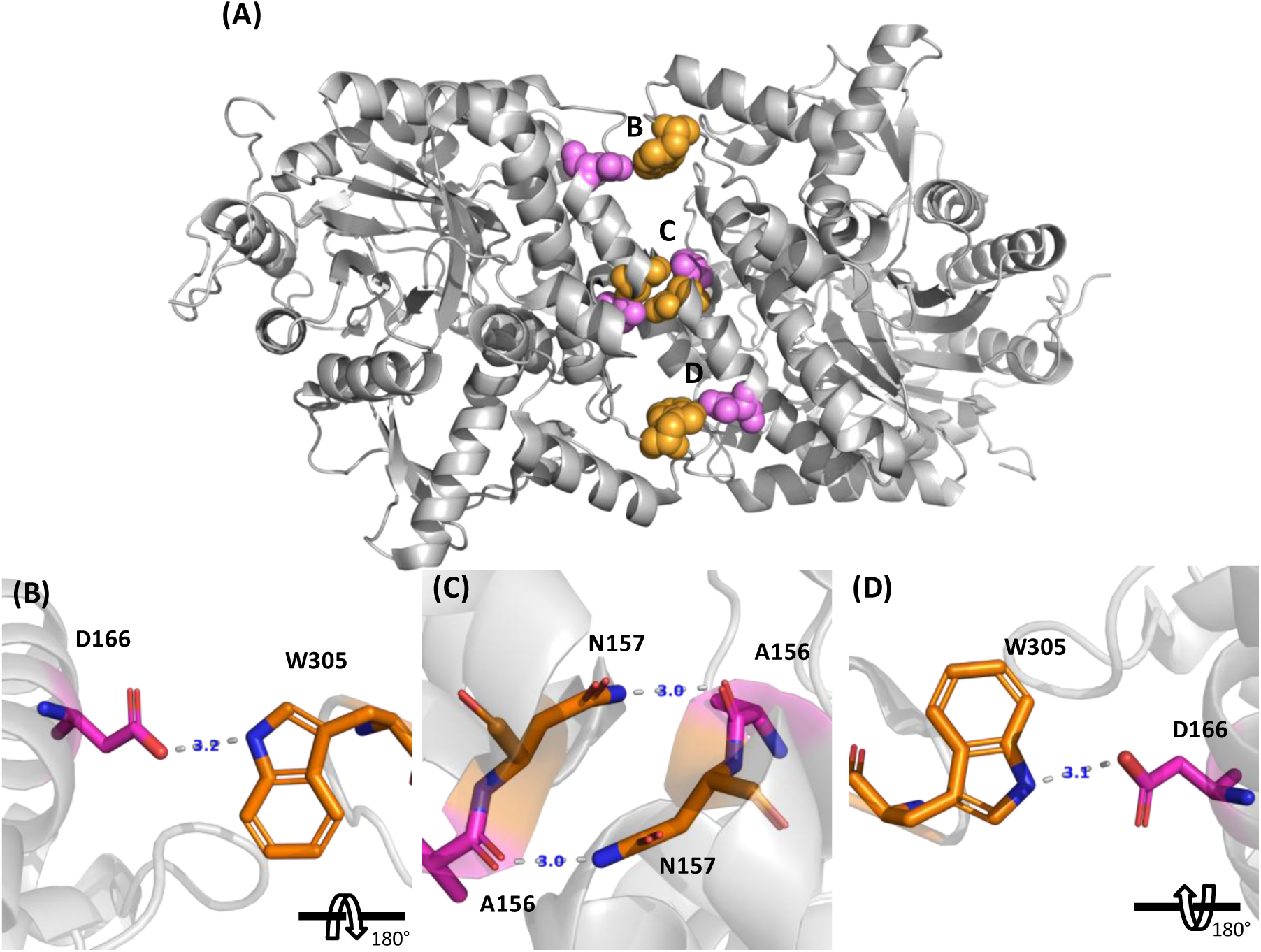
Hydrogen bonds at the Sfβgly homodimer interface (PDB 5CG0). **(A)** Residues in pink are acceptors, while in orange are donors. **(B)**, **(C)** and **(D)** panels detail the hydrogen bonds (dashed lines) and their positions at the interface. Distances (in blue) are in Å. The letters are the same as in panel **(A)**: B and D indicate interactions at the interface extremities and C interactions at the center.

The hydrogen bond between N157 and A156 corresponds to 2.4 *k*J/mol, while the bond involving D166 and W305 corresponds to 5.9 *k*J/mol (Table 2). These values are typical of hydrogen bonds in proteins [28]. Moreover, they indicate that, collectively, the four hydrogen bonds contribute 16.6 *k*J/mol (2 x 2.4 + 2 x 5.9) to the stabilization of the Sfβgly homodimer.

#### 2.3.3 Mutational effects on the enzymatic activity

Finally, considering that dimerization affects the Sfβgly catalytic activity, the mutational effect on the enzyme kinetic parameters was evaluated (Table 3).

**Table 3.** Enzyme kinetics parameters for the wild-type and mutant Sfβgly.

| Sfβgly | $k_{cat}$ ($\text{min}^{-1}$) | $K_m$ (mM) | $k_{cat}/K_m$<br>($\text{min}^{-1} \cdot \text{mM}^{-1}$) |
| --- | --- | --- | --- |
| wild-type | $143 \pm 2$ | $1.1 \pm 0.1$ | 122.2 |
| N112S | $140 \pm 4$ | $1.2 \pm 0.2$ | 109.3 |
| N157S* | $66 \pm 2$ | $1.1 \pm 0.2$ | 60 |
| D166S* | $80 \pm 1$ | $2.2 \pm 0.2$ | 35.7 |
| M210A | $116 \pm 2$ | $1.1 \pm 0.2$ | 99.1 |
| L214A | $167 \pm 7$ | $1.9 \pm 0.4$ | 84.8 |
| Y303A | $73 \pm 3$ | $2.6 \pm 0.5$ | 27.2 |

As expected those mutations reduced the Sfβgly activity, an effect credited to the decrease in the homodimer formation, *i.e. K*_D_ increment (Table 2). Nevertheless, as the replacement of apolar residues had a more intense effect on the monomer affinity than the mutational disruption of the hydrogen bonds (Table 2), their effect on the Sfβgly activity was separately analyzed within these two classes. Indeed, the replacement of residues involved in the hydrophobic effect led to a gradual linear decrease in *k*_cat_/*K*_m_ as *K*_D_ increased (Figure 6), whereas mutations in residues involved in hydrogen bond formation resulted in an abrupt decline. As fewer mutants are available in the second case, it is not viable to draw a *k*_cat_/*K*_m_ and *K*_D_ correlation as in the former. Anyway, the hydrogen bond disruption seems to have a supplementary damaging effect on the activity, which goes beyond the plain reduction in the concentration of the homodimer.

**Figure 6.**
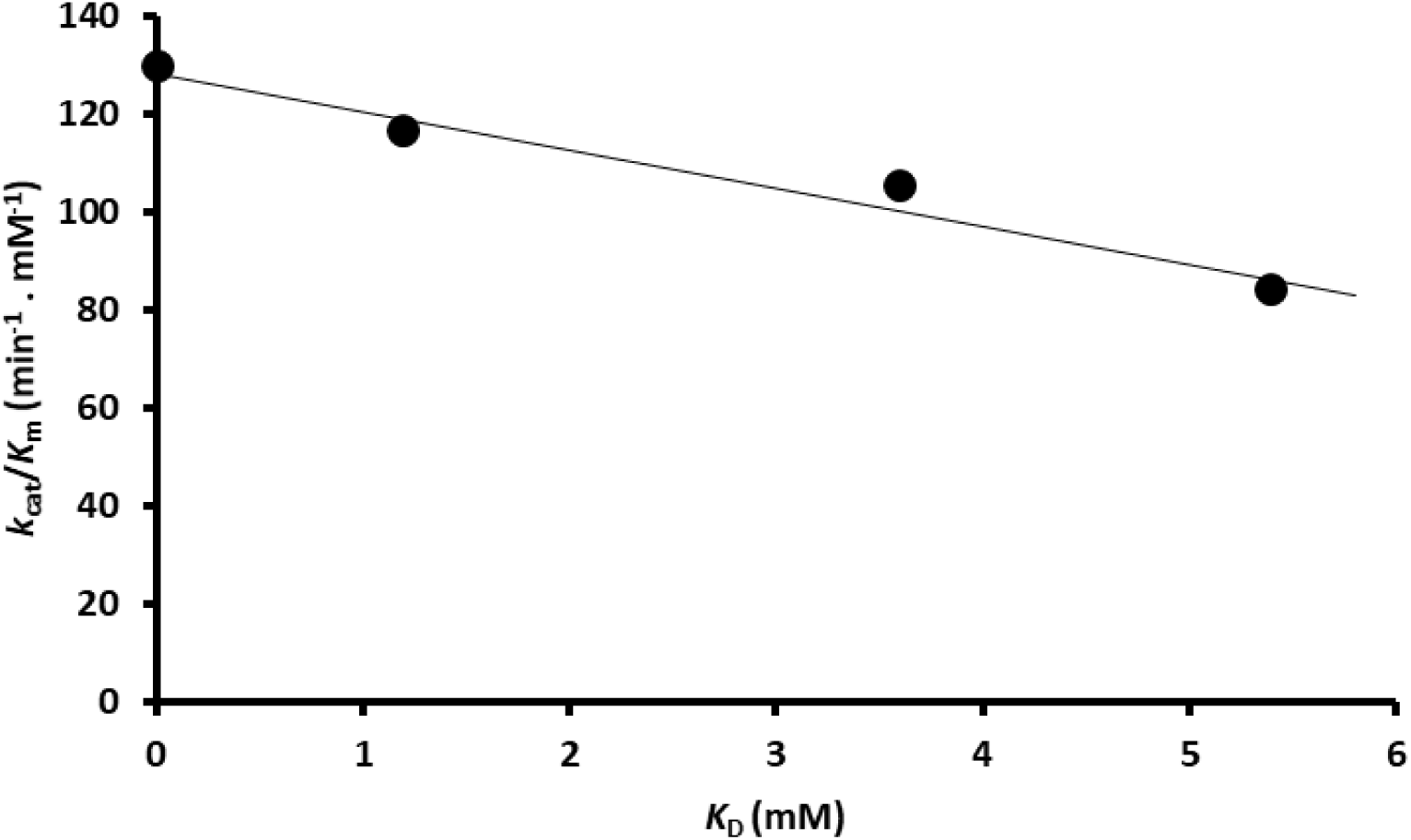
Catalytic efficiency (*k*_cat_/*K*_m_) and *K*_D_ of Sfβgly for mutations of apolar residues involved in the hydrophobic effect. Dots represent the wild-type Sfβgly, N112S, M210A, and L214A mutants, respectively (Tables 2 and 3).

## 3. Discussion

The crystallographic asymmetric unit does not necessarily reflect the biological oligomerization state of a protein, so a crucial task is to distinguish biologically significant protein interfaces from nonbiological interfaces in a crystal [29, 30]. PDBePISA server, which provides accurate predictions in over 80% of cases [1], was used to estimate the oligomerization state of 88 GH1 β-Glucosidases crystallographic structures. Such analysis showed that 47% of those enzymes are monomeric in solution, while 30% are dimers, 2% are trimers, 14% are tetramers, 5% are hexamers, and 2% are octamers. These data are consistent with previous arguments that at least 50% of hydrolases form oligomers [14]. Such oligomer predominance reinforces the relevance of the detection and characterization of the Sfβgly homodimerization in solution, under biologically relevant conditions.

Some proteins are predominantly found in their oligomeric state, exhibiting *K*_D_ in the nanomolar range [8]. In contrast, other proteins exhibit a weaker tendency to associate and are classified as transient. In this case, they have higher *K*_D_, ranging from micromolar to millimolar [31, 32], which is the scale for Sfβgly dimer. Regarding the analysis of the Sfβgly dimerization at different concentrations of the phosphate buffer, the *K*_D_ was here shown to vary between 7 to 135 µM, depending on the ionic concentration.

The literature refers to those dimers, whose *K*_D_ ranges from micromolar to millimolar scale, as weak dimers and also states that most of them burry between 1200 to 2000 Å^2^ at their interfaces [33], recapping that the dimeric interface of Sfβgly buries 905 Å^2^. A smaller interface could imply fewer non-covalent interactions and a smaller contribution of the hydrophobic effect [31], which probably explains the low stability of Sfβgly homodimers.

Interfacial residues N112, M210, L214, and Y303 contribute only a small fraction of the solvent-accessible surface area of the buried interface within each monomer (9%, 3%, 5%, and 4%, respectively). However, it should be considered that their replacement for smaller residues not only removed the surface area at those restricted points, but also exposed the surface of the residues with which they interact at the interface. N112 interacted with Y303, which was also one of the mutated residues. Additionally, M210 contacted M110, and L214 interacted with L159 and W163. Together, the mutated residues and their interacting counterparts account for 16%, 12%, and 19% of the buried surface area of the interface within each monomer, respectively. Additionally, it should be noted that these pairs are duplicated at the interface, so each one corresponds to approximately 30% of the solvent-excluded surface area of the interface (32%, 24%, and 38%, respectively).

In summary, these sets of residues involved in the hydrophobic effect at the interface contribute similarly, with approximately 15 *k*J/mol each one (Table 2), to the affinity between the monomers of the Sfβgly dimer. These experimental values are close to those expected based on the contribution of 54.4 J/mol per Å² of buried apolar surface area at protein interfaces [34], which would correspond to 15, 11, and 18 *k*J/mol for N112 - Y303, M210 - M110, and L214 - L159/W163, respectively.

As these residue sets correspond to approximately 30% of the solvent-accessible surface area buried in the dimer, but they account for 50% of the affinity between monomers, the removal of about one-third of the interface surface area from the solvent appears to be sufficient to stabilize the interaction between monomers, compelling them to the association. It also reinforces the hypothesis, based on the ionic concentration effect on the *K*_D_ (Figure 2), that the hydrophobic effect is crucial for stabilizing the Sfβgly homodimer.

Remarkably the contribution from the hydrophobic effect of just one of the residues mentioned above (N112, M210, L214, and Y303; ∼15 *k*J/mol) is similar to the contribution of the complete set of the interfacial hydrogen bonds to the stabilization of the Sfβgly homodimer, which is 16.6 *k*J/mol (Table 2). This observation points to a specific role for the hydrogen bonds. Indeed, they are distributed over the interface in a pattern that likely ensures specificity in the protein-protein interaction (Figure 5).

Indeed, as commented above, slightly more than 30% of the interface surface of each monomer interacting *via* the hydrophobic effect suffices for an inevitable dimerization. It would be precisely here, appreciating that the monomers have plenty of apolar interface area for misplaced binding, that hydrogen bond forming residues are revealed critical for ensuring specificity, preventing the monomers from binding in an unviable orientation.

The position of hydrogen bonds at the interface and the corresponding mutational effect of their perturbation provide additional support to this hypothesis. The D166S mutation reduces the monomer affinity to 5.9 *k*J/mol, while the N157S mutation results in a decrease of 2.4 *k*J/mol. This difference may be related to the positioning of these residues within the dimer interface. The D166 residues are located at both extremities of the interface, while the N157 residues are situated more centrally. So, the hydrogen bonds at the extremities, between D166 and W305, seem to play a more important role in maintaining homodimer stability. Disrupting these bonds at the interface extremity would have a greater impact on stability compared to disrupting hydrogen bonds near the center because the wobbly edges would expose hydrophobic patches to the solvent. In short, hydrogen bonds involving D166 and W305 would seal the interface edges.

Interestingly, the mutational effects on the Sfβgly activity may be additional evidence of the differential role of the hydrophobic and hydrogen bonding residues in the homodimer interface. So, assuming that the hydrophobic effect only depends on the area of the apolar surface removed from the solvent with no particular requisite on its positioning, any change in the apolar area would equally affect the dimer-to-monomer ratio. Hence, the continuous decrease of the *k*_cat_/*K*_m_ resulting from the decrement in the monomer affinity due to the loss of apolar surface in their interfaces might be simply related to the reduction of dimer concentration (Figure 6).

Conversely, the *k*_cat_/*K*_m_ abrupt decline observed in the interfacial hydrogen bonds disruption hint at an extra mutational effect in addition to the simple modification of the homodimer to monomer ratio (Table 3). One could speculate that in the dimer bearing a hydrogen bond rupture at a crucial point of its interface, the monomers might remain associated, but subtly misaligned. Hence, the alteration of their relative positions could prevent one monomer from accurately fit in the other, reducing their activity even in the dimeric state.

Finally, these observations about the hydrogen bond distribution and also the dimer instability when more than 30% of the interface becomes exposed to the solvent can be combined in a picture of the homodimer assembly and disassembling. The dimerization would involve two Sfβgly monomers approaching due to the hydrophobic effect. When more than 30% of their interface surfaces were hidden from the solvent, the homodimer would be stable. However, the ligation would progress up to the point that D166 and N157 were properly aligned for the hydrogen bond formation in the center of the interface. Such anchorage point would be reinforced by the burial of the hydrophobic residues around. Finally, the hydrogen bonds in the interface extremities, involving D166 and W305, would zip the association. Moving on, the continuous dynamics of the homodimer would eventually lead to a temporary separation at the peripheries of the interface, *i.e.* the hydrogen bond between D166 and W305 could be weakened or ruptured. Such temporary departure would open a breach to the water getting in. If that gap advanced beyond 70% of the interface surface then the dimer would collapse.

## 4. Materials and Methods

### 4.1 Site-directed mutagenesis

Site-directed mutagenesis experiments were performed using QuikChange Lightning Site-Directed Mutagenesis Kit (Agilent Technologies, Santa Clara, CA, USA) according to manufacturer instructions. The pET-46 EK-LIC vector (Merck Millipore, Billerica, MA) containing the insert coding for Sfβgly (pET46-Sfβgly) was available in our group and served as the template for producing the mutants.

### 4.2 Expression and purification of recombinant proteins in bacteria

The WT Sfβgly and mutants were expressed and purified according to the method described previously [20]. Following purification, the wild-type and mutant Sfβgly samples underwent buffer exchange using PD Minitrap G-25 columns (Cytiva, Marlborough, MA, USA). The final samples were stored in a 100 mM phosphate buffer pH 6 (P100) at 4°C. Sample purity was evaluated through SDS-PAGE [35] and protein concentration was determined using the bicinchoninic acid (BCA) assay [36].

To perform SEC chromatography of wild-type Sfβgly in buffers other than P100 the stored samples were exchanged to 75 mM (P75), 50 mM (P50), 25 mM (P25), or 5 mM (P5) phosphate buffer pH 6 using PD Minitrap G-25 columns.

### 4.3 Size exclusion chromatography (SEC)

The SEC was conducted using an ÄKTA FPLC (GE HealthCare) system. Two Superdex 200 HR10/300 GL (GE HealthCare) columns were coupled in tandem and equilibrated with appropriate phosphate buffer pH 6. The columns were placed on crushed ice to enhance peak resolution. In that condition, the system pressure stabilized at 1.1 MPa with a flow rate of 0.25 ml/min. The wild-type and mutant Sfβgly samples were loaded into the system using a 100 μl sample loop. Data were collected and processed using the UNICORN software version 5.11 (GE HealthCare). The area of the chromatographic peaks was used to estimate the relative abundance of monomers and dimers, which combined with the total protein concentration, resulted in the monomer and dimer concentration. Those data were used in the *K*_D_ determination as following. Assuming a simple equilibrium between dimer (D) and monomer (M) Sfβgly

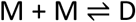

the dissociation constant (*K*_D_) is expressed as

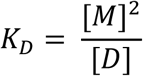

which can be rearranged to

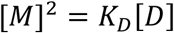

indicating that a plot of [M]^2^ *versus* [D] should be a line and its slope would correspond to the *K*_D_

### 4.4 SEC-MALS

To determine the molar mass of the peaks observed in wild-type Sfβgly samples in P100, a SEC system as described above was connected to a miniDAWN TREOS multi-angle static light scattering (MALS) and an Optilab T-rEX detector (Wyatt Technology Corporation, Santa Barbara CA). Experimental conditions were the same as those previously described. Data were collected and analyzed using the ASTRA software version 7.1.1.3 (Wyatt Technologies).

### 4.5 Enzyme activity of wild-type and mutant Sfβgly

The initial rate (*v*_0_) of hydrolysis of at least 10 different concentrations of the substrate *p*-nitrophenyl-β-Dglucopyranoside (NPβglc) prepared in P100 buffer was used to determine the enzyme kinetic parameters (*k*_cat_ and *K*_m_). The diluted enzyme (50 μl) was mixed with the substrate (50 μl) in a 96-well plate while on ice. The enzymatic reaction began when the assay plates were moved to a 30**°**C warm bath. To determine the initial rate, reactions at each substrate concentration were performed at four times intervals (5, 10, 15, and 20 min) to monitor product accumulation as a function of time. The reactions were stopped by adding 0.5 M Na_2_CO_3_ (100 μl). Absorbance at 415 nm was measured using an Elx800 microplate reader (Biotek) and converted to moles of the released product (4-nitrophenol) using a linear calibration curve. The product (in nmol) was plotted against the respective time intervals (5, 10, 15, and 20 min), and the *v*₀ (nmol/min) was determined from the slope of the linear plots. The substrate concentration and their corresponding *v*₀ were fitted to the Michaelis–Menten equation to determine the kinetic parameter (*K*_m_, *k*_cat_, and *k*_cat_/*K*_m_). These assays for kinetic parameter determination were repeated three independent times.

### 4.6 Circular dichroism (CD) experiments

The wild-type and mutant Sfβgly secondary structures were evaluated by collecting circular dichroism (CD) spectra on a Jasco-815 spectropolarimeter. Samples consisted of 5 μM of protein in P100. Spectra were collected over a range of 260 nm to 200 nm with a scan speed of 20 nm/min and 3 accumulations to increase the signal-to-noise ratio. The molar ellipticity was calculated as previously described [37].

## Author Contributions

**Rafael S. Chagas:** conceptualization; formal analysis; investigation; writing original draft; writing review and editing. **Sandro R. Marana:** conceptualization; funding acquisition; resources; formal analysis; investigation; writing - original draft; writing - review and editing.

## Supplementary Material

**Supplementary Figure 1.**
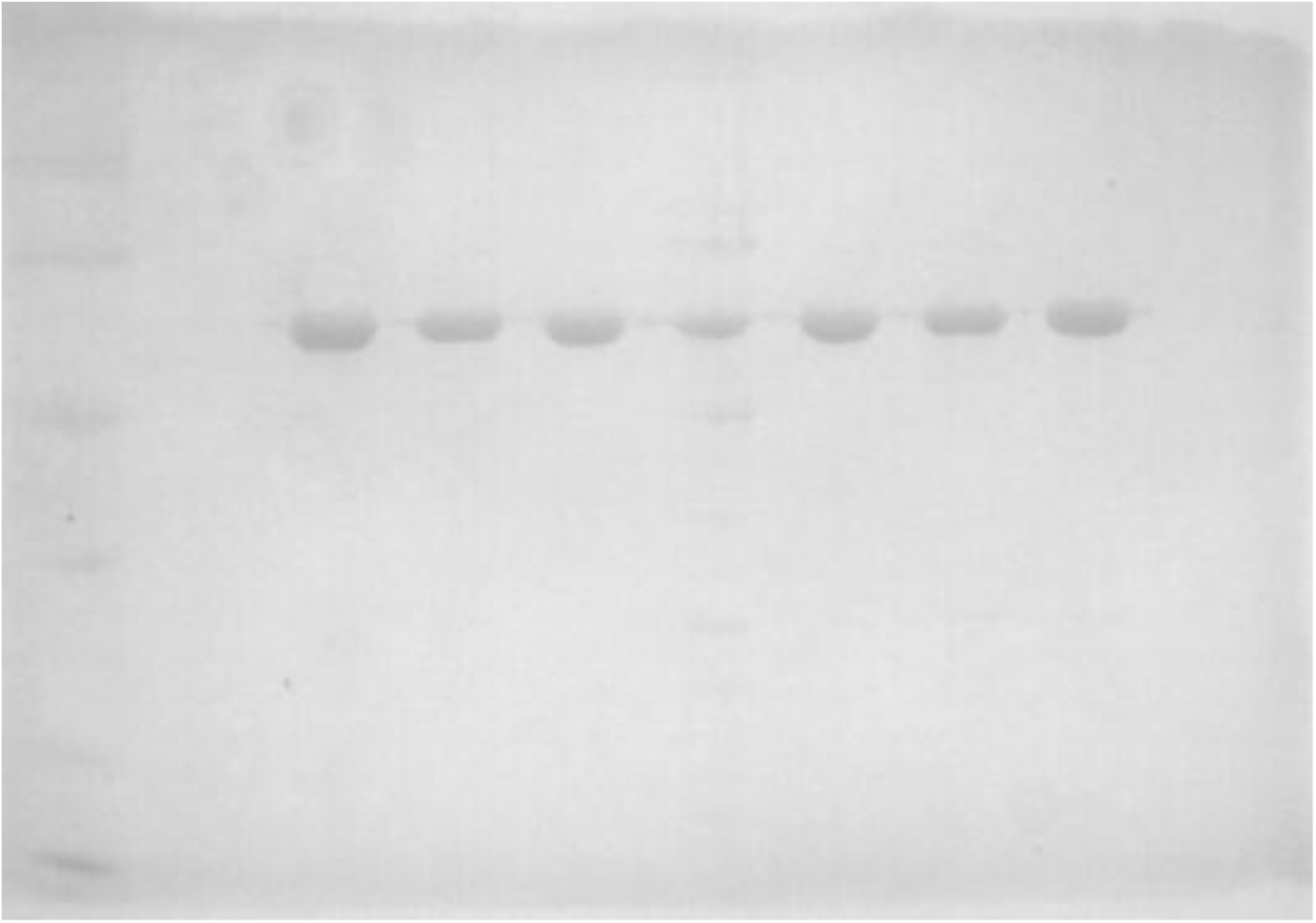
SDS-PAGE of the purified recombinant wild-type and mutant Sfβgly. **1** – Wild-type Sfβgly; Mutant Sfβgly are: **2** – N112S; **3** – N157S; **4** – D166S; **5** – M210A; **6** – L214A; **7** – Y303A.

**Supplementary Figure 2.**
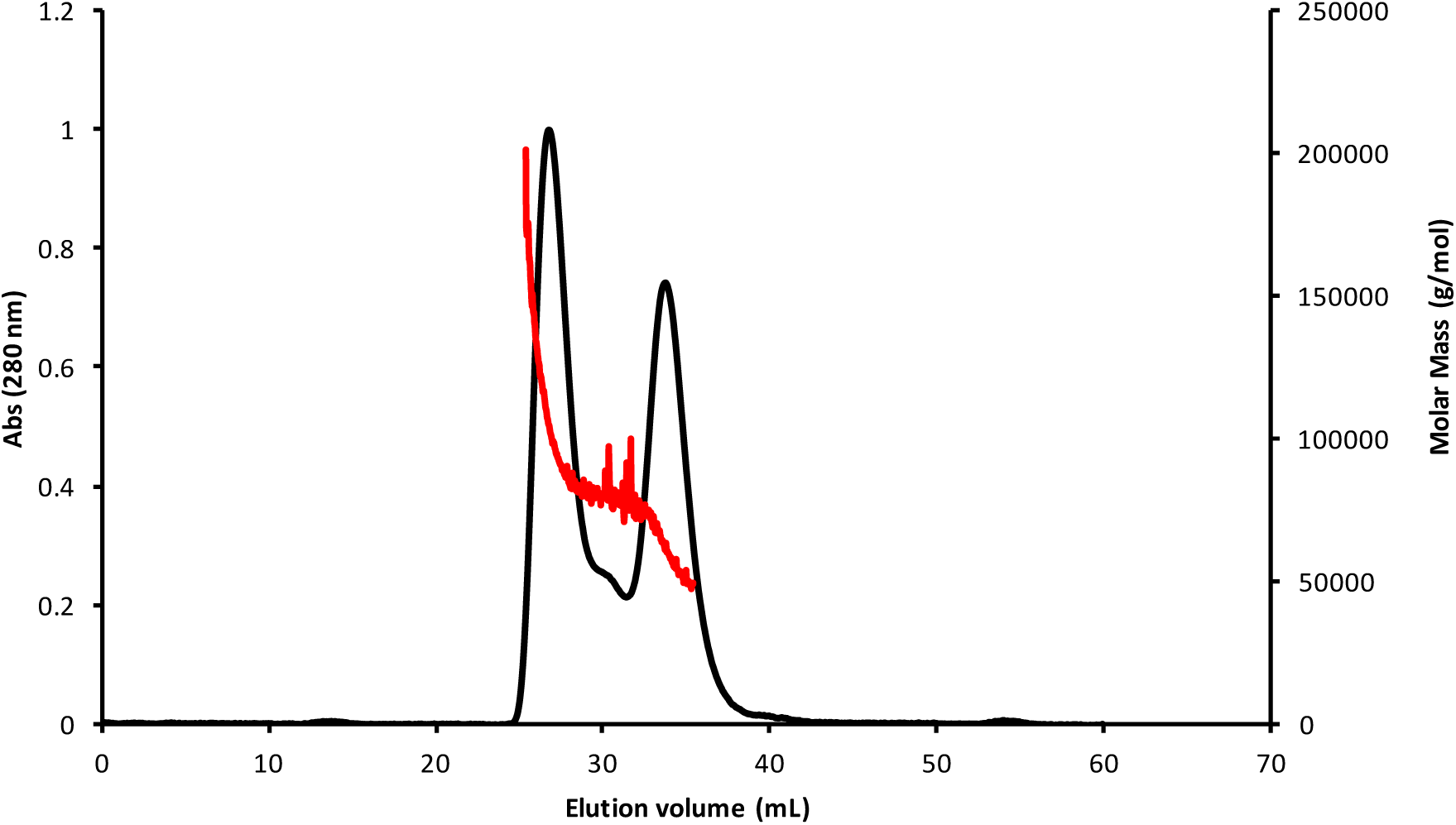
Analysis of the purified wild-type Sfβgly in size exclusion chromatography coupled to a multiple angle light scattering detector (SEC-MALS). Two different oligomerization states (dimer and monomer were observed when μM Sfβgly was analyzed at 5°C.

**Supplementary Figure 3.**
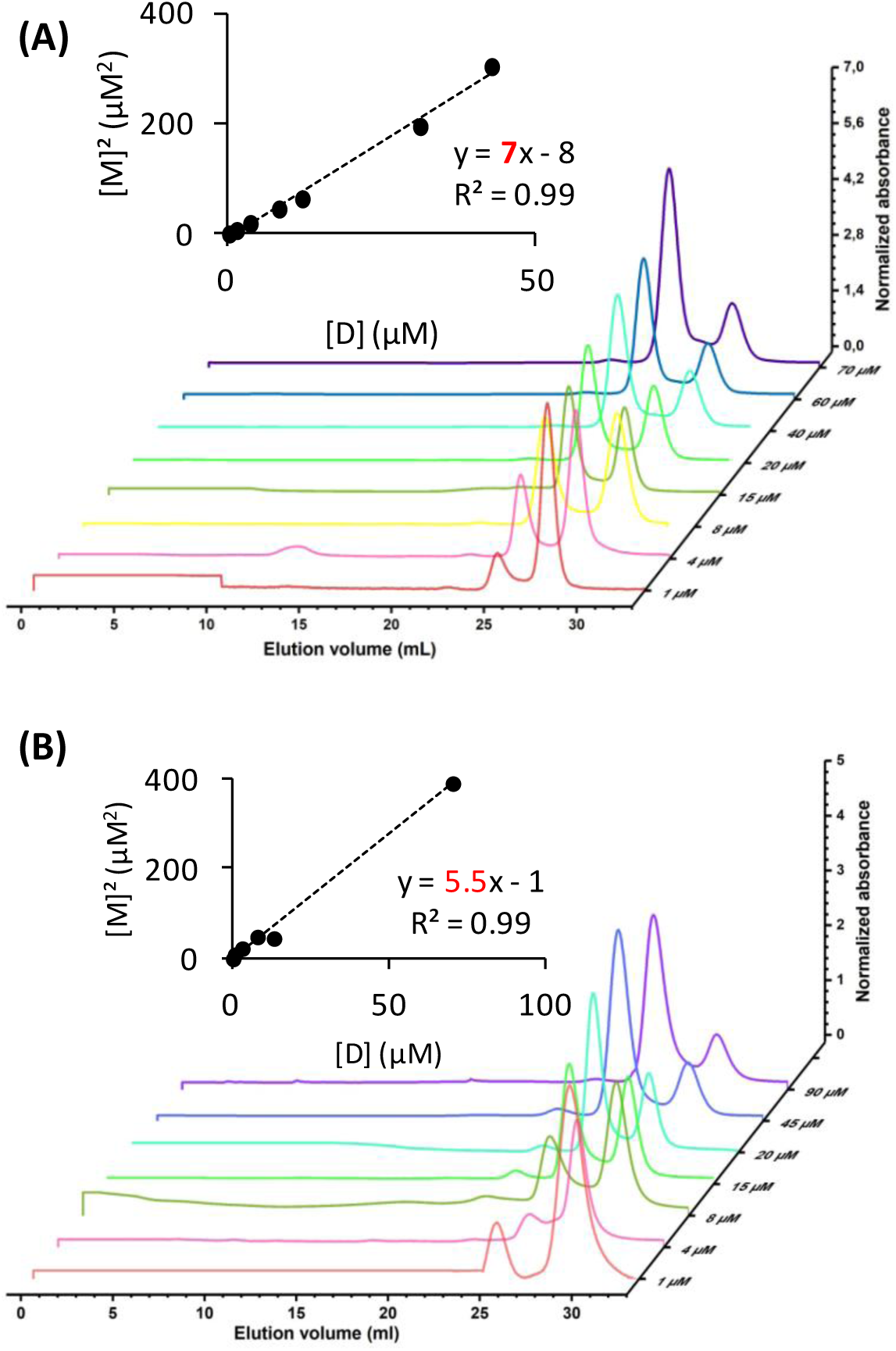
Determination of the dissociation constant (*K*_D_) of the Sfβgly dimer in two independent experiments. (**A**) Size exclusion chromatography (SEC) of the purified wild-type Sfβgl over di erent ini al protein concentra ons at C in 100 mM Phosphate buffer pH 6 (P100). (**B**) Plots for estimation of the *K*_D_ for the wild-type Sfβgl. Details in Materials and Methods.

**Supplementary Figure 4.**
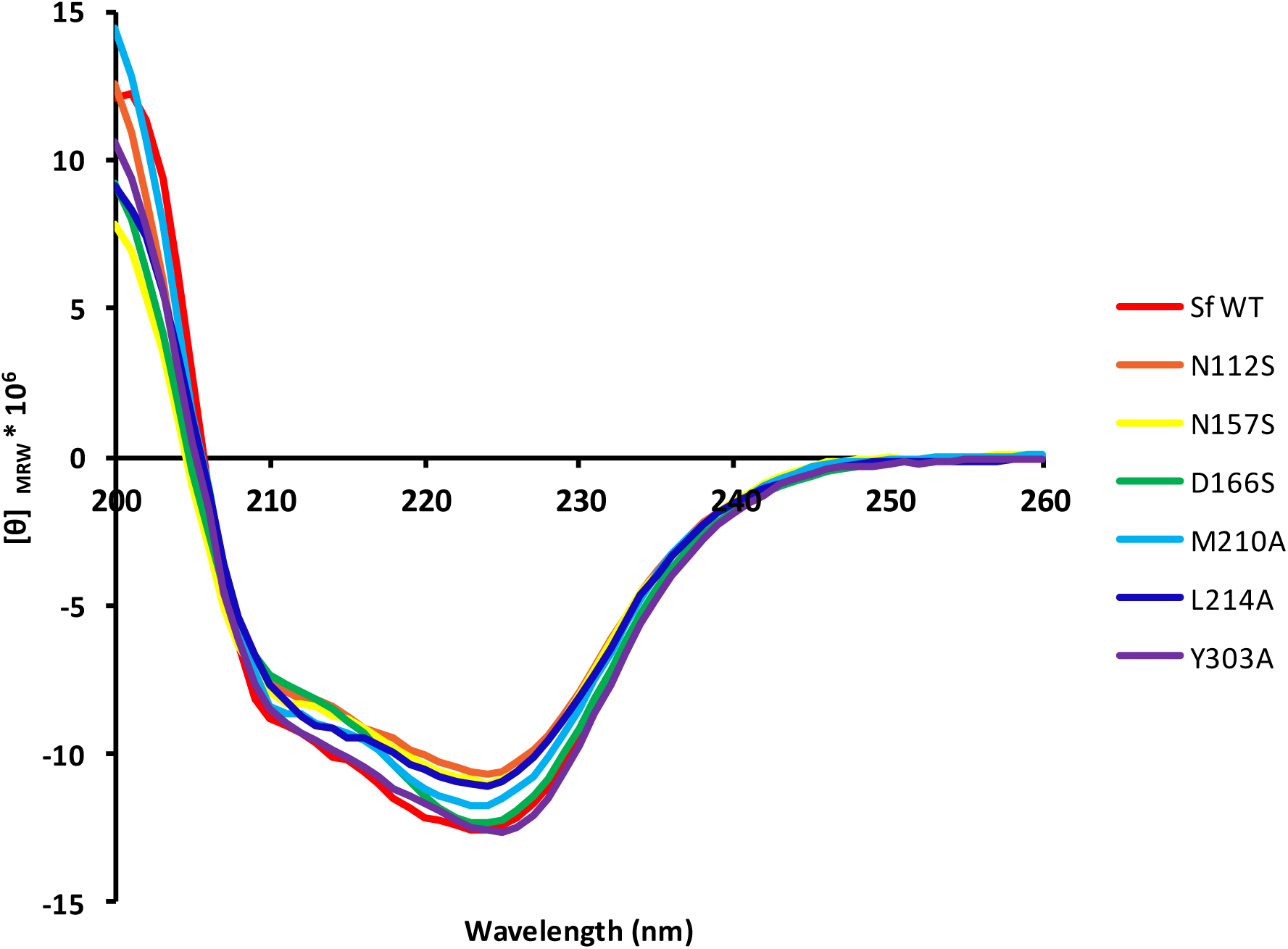
Circular dichroism spectra of the wild-type and mutant Sfβgly. Different colors indicate different mutants.

**Supplementary Figure 5.**
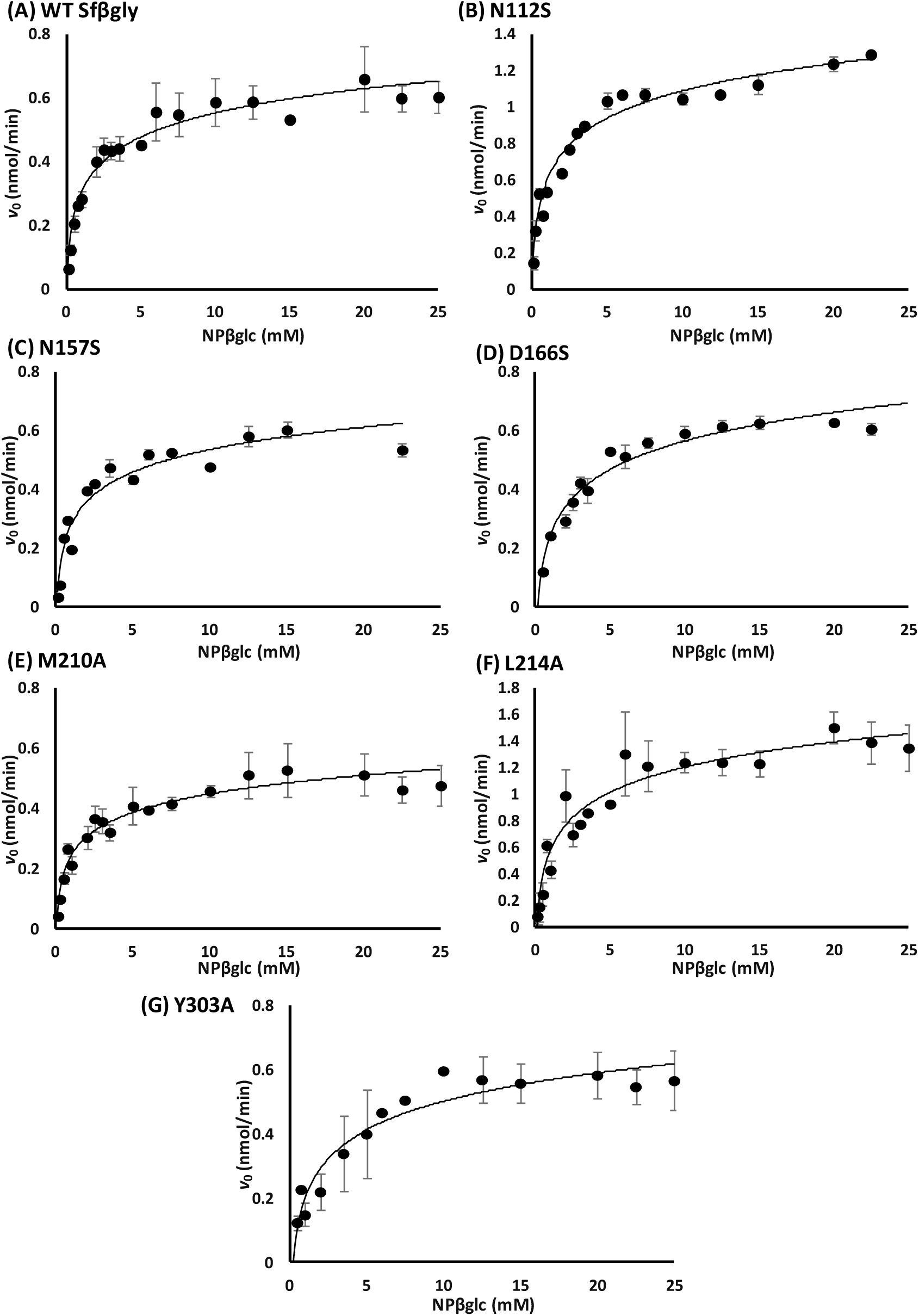
Effect of NPβglc concentration on the activity of the wild-type and mutant Sfβgly. (**A**) wild-type Sfβgly, 0.0025 µg/µL; **(B)** N112S, 0.005 µg/µL; **(C)** N157S, 0.005 µg/µL; **(D)** D166S, 0.005 µg/µL; **(E)** M210A, 0.0025 µg/µL; **(F)** L214A, 0.005 µg/µL; **(G)** Y303A, 0.005 µg/µL.

